# Confidence Judgments Reflect the Standard Error of Noisy Evidence Samples Across Domains

**DOI:** 10.64898/2026.04.20.719573

**Authors:** Rebecca K. West, David K. Sewell, Benjamin Sheibehenne

**Affiliations:** Technical University of Munich, Germany; University of Queensland, Australia; Karlsruhe Institute of Technology, Germany

## Abstract

Confidence judgments play a critical role in guiding behavior by shaping information-seeking, learning, and decision strategies. These functions are most effective when confidence is well calibrated, that is, when subjective uncertainty aligns with the objective uncertainty in the presented evidence. Motivated by this, we investigated how people form confidence judgments from noisy samples of information, and whether they use statistically grounded strategies or rely on heuristics. Participants performed two categorization tasks, one with visual orientation stimuli and one with number stimuli. In each task, participants saw sequentially presented observations and made a decision about the generating category and simultaneously reported their confidence in that decision. We independently manipulated the number of observations and standard deviation of the sample to assess whether confidence reflected an integrated estimate of both sources of statistical uncertainty. Behaviorally, confidence and accuracy both increased with larger sample sizes and lower variability. Furthermore, confidence and accuracy were equivalent in samples matched for standard error, suggesting that participants relied on a statistically grounded strategy. Computational modeling further supported this interpretation: a model that scaled confidence according to the standard error of the sample mean provided the best fit to the data, outperforming more heuristic and Bayesian alternatives. This pattern generalized across the orientation and number tasks, suggesting a domain-general strategy for uncertainty estimation. Together, these findings demonstrate that people use structured, statistically grounded strategies to compute their confidence, supporting well-calibrated decision-making even in the absence of full Bayesian inference.

## Introduction

In the real world, people often have to make decisions based on limited and noisy information. For example, evaluating the quality of a product from a handful of reviews or estimating the volatility of a stock from past market activity requires combining multiple uncertain observations into a single judgment. As part of this process, people also form a sense of how certain they are that their judgment is correct. This subjective sense of certainty, or *decision confidence*, is a form of metacognition because it involves reflecting on the quality of one’s own decisions. But how do people compute confidence from noisy evidence samples?

This question matters because confidence is not just a passive outcome of the decision process but plays an active role in guiding behavior. Confidence judgments serve as an internal signal that help regulate decision strategies, shaping how much evidence people gather before making a choice (Balsdon et al., 2020; van den Berg et al., 2016), whether additional information is sought after an initial decision (Desender et al., 2018), and how people learn from past decisions in the absence of external feedback (Cortese et al., 2020; Daniel & Pollmann, 2012; Guggenmos et al., 2016; Lovelace, 1984; Son & Metcalfe, 2000; Thiede & Dunlosky, 1999). However, confidence judgments are only useful for guiding behavior to the extent that they are well-calibrated. That is, they must accurately reflect the *diagnostic value* of the underlying evidence. A central question in this study, therefore, is *how* people compute confidence from uncertain evidence and whether confidence judgments accurately track the properties of the evidence on which they are based.

From a Bayesian perspective, confidence corresponds to the posterior probability that a decision is correct given observed evidence and prior knowledge (Fleming & Daw, 2017; Meyniel et al., 2015; Pouget et al., 2016). Bayesian models provide a principled account of how confidence should vary with the diagnostic value of evidence, and they have been shown to capture key aspects of human confidence judgments in some contexts (Aitchison et al., 2015; Hangya et al., 2016; Li & Ma, 2020; Navajas et al., 2017; Sanders et al., 2016). However, other empirical findings suggest that people do not always behave in accordance with Bayesian principles. For example, they may discount or overlook evidence that contradicts their chosen option (Maniscalco & Lau, 2016; Zylberberg et al., 2012) or they may be influenced by decision-irrelevant sources of noise (Spence et al., 2016). These deviations raise the possibility that people rely on simpler, heuristic strategies when computing confidence.

Indeed, alternative theoretical accounts propose that subjective confidence reflects approximate or heuristic evaluations of evidence rather than fully optimal probabilistic computations. For instance, Griffin and Tversky (1992) distinguish between the strength of evidence (i.e., the balance of evidence for one option over another) and the weight of evidence (i.e., the reliability or diagnosticity of that evidence). Their findings suggest that people often focus disproportionally on strength while neglecting weight, leading to overconfidence when strong evidence is based on a small or unreliable sample, and under confidence when weak evidence is based on highly reliable information (Tversky & Kahneman, 1971, 1974). These findings highlight the need to better understand whether and how people incorporate the diagnostic value of evidence into their confidence judgments, especially in situations where that evidence is uncertain or noisy.

A particularly relevant form of uncertainty involves uncertainty in the sample mean, that is, how precisely one can estimate a population value based on a finite set of noisy observations. This uncertainty can be jointly determined by two key features of a sample: sample size and sample variability. Larger samples generally provide more reliable estimates by reducing uncertainty in the sample mean, whereas greater variability in the observations increases that uncertainty. Together, these factors determine the standard error of the sample mean, a statistical measure that quantifies the expected precision of an estimate. Crucially, the standard error inherently reflects the trade-off between sample size and sample variability where a small, low-variability sample might yield a more precise estimate than a large, highly variable one. Despite the relevance of this trade-off, few studies have manipulated sample size and variability orthogonally to test how they jointly influence confidence judgments.

One exception is a recent study by Olschewski and Scheibehenne (2024), who found that when participants estimated the mean of number samples matched for standard error, their responses were less accurate and less confident for large, high-variance samples than for small, low-variance samples. The authors attribute this effect to increased cognitive imprecision when integrating numerous noisy samples, suggesting that confidence judgments may reflect processing limitations rather than solely statistical uncertainty in the stimuli. However, open questions remain about the generalizability of these findings to other domains of decision-making and response formats.

More broadly, it is unclear whether the mechanisms underlying confidence judgments generalize across different domains of decision-making or whether they are specific to particular stimulus domains (Faivre et al., 2018; Mazancieux et al., 2020; Navajas et al., 2017; Rouault et al., 2018; West et al., 2023). Several accounts propose that confidence reflects a common metacognitive process that operates across different types of decisions (Fleming & Daw, 2017; Yeung & Summerfield, 2012), potentially supported by a domain-general neural system (Morales et al., 2018). Behavioral studies have provided partial support for this view, showing positive correlations in metacognitive sensitivity across perceptual and cognitive tasks (de Gardelle & Mamassian, 2014; Faivre et al., 2018; Rouault et al., 2018). However, other findings suggest important domain-specific influences. For example, metacognitive efficiency appears to vary across tasks (Mazancieux et al., 2020), and confidence judgments can be differentially affected by the format and modality of evidence (Ais et al., 2016). Moreover, it remains unclear whether quantitative manipulations of sample uncertainty, such as changes in sample size and variability, influence confidence similarly across domains. Addressing this question is crucial for testing the extent to which confidence reflects a general computation of uncertainty versus one that is tuned to the statistical structure of specific types of evidence.

To address these gaps, the present study systematically manipulated sample size and sample variability in a probabilistic categorization task across two stimulus domains: oriented gratings and number stimuli. In an online experiment, participants (*N* = 85) viewed a sequence of *observations*, referred to collectively as a *sample*, and were instructed to use the mean of the presented sample to make a judgment about which of two categories (category 1 or category 2) it had been drawn from. Participants made a simultaneous judgment about the category and their confidence in that decision, indicating their response on a continuous scale from 0% to 100% category 2. We varied both the number of observations in the sample (4 or 16 observations) and the variance of the sample (small, medium, large) to investigate the effect of both sources of uncertainty on decision accuracy and confidence. We hypothesized that both accuracy and confidence would increase with larger sample sizes and with lower variability, as both factors reduce uncertainty in the sample mean.

Critically, we also included matched standard error conditions, pairs of trials where a small, low-variance sample and a large, high-variance sample had equivalent statistical uncertainty, to assess whether participants’ decisions and confidence reflected the trade-off between sample size and sample variability. We hypothesized that if confidence judgments are based on statistically optimal computations, participants should exhibit comparable accuracy and confidence across these matched conditions. Conversely, if confidence arises from heuristic strategies, such as overweighting sample size or variability, or is shaped more by perceived uncertainty than statistical uncertainty (Olschewski & Scheibehenne, 2024), we would observe differences in accuracy and confidence across the matched conditions. Finally, if confidence is computed using domain-general mechanisms, we would expect the observed patterns to generalize across both the orientation and number tasks.

To formally test whether participants’ judgments reflected statistically optimal integration of uncertainty, we fit a series of computational models. These models varied in complexity, ranging from those that relied solely on the sample mean, to models that incorporated sample size and variance as separate or interacting factors, to a model that computed statistical uncertainty using the sample’s standard error. We also compared these models to a Bayesian observer model (Adler & Ma, 2018; Aitchison et al., 2015; West et al., 2023) that computed a posterior probability ratio using all sample statistics and prior category information. This modeling framework allowed us to assess whether participants’ confidence judgments were better explained by optimal inference or heuristic approximations, and whether the same computational strategies applied across domains.

Consistent with a statistically grounded strategy, we found that when samples were matched in standard error, participants’ accuracy and confidence did not differ. Model comparisons further supported this pattern: the best-fitting model assumed that participants accounted for uncertainty in the sample mean using the sample’s standard error. This model outperformed alternatives that assumed no sensitivity to uncertainty, treated sample size or variability in isolation, or implemented a full Bayesian decision rule. The model also provided the best fit for both the orientation and number task, suggesting that participants used a consistent strategy for integrating uncertainty in both domains. Overall, our findings offer new insights into how people form confidence judgments under uncertainty across different decision contexts and contribute to broader debates about the nature and limits of metacognition.

## Method

### Overview

On each trial, participants viewed a sample of stimuli on a computer screen and made a probability judgment about its category membership, estimating whether it had been generated by ‘category 1’ or ‘category 2’. Observations were presented sequentially, and we manipulated the sample size (4 or 16 observations), standard deviation (small, medium, or large), and domain (oriented gratings or number values). Participants used a slider scale to indicate their probability judgment and were incentivized using a Brier scoring rule.

### Participants

We conducted an a-priori power analysis using G*Power, targeting 80% power and an alpha level of 0.05 to detect small to medium effect sizes (Cohen’s f^2^ = 0.15). The analysis, based on linear multiple regression with fixed effects for sample mean, sample SD, sample size, and an intercept term, suggested a required sample size of 85 participants.

123 participants were recruited through Prolific and reimbursed for their time. All participants reported normal (or corrected-to-normal) vision and English comprehension. Participants completed the task online using a desktop or laptop computer; mobile devices were not allowed. After applying exclusion criteria (see Results), data from 85 participants remained, consistent with the power analysis target.

### Experimental Procedure

As shown in **Figure 1**, a fixation cross was presented in the center of a mid-grey background for 500 ms at the start of each trial. Following fixation, participants viewed a sample of 4 or 16 stimuli, with each observation presented sequentially for 400 ms, separated by a blank 100 ms inter-stimulus interval. In the orientation task, stimuli were oriented sinusoidal gratings. In the number task, stimuli were numbers. On each trial, the sample was drawn from one of two categories, each defined by a Gaussian distribution with different means but the same variance. In the orientation task, the category means were oriented left (*μ*_1_ *=* −5°) or right (*μ*_2_ *=* 5°) of horizontal; in the number task, distributions were shifted such that they were centered above (*μ*_1_ *=* 195) or below (*μ*_2_ *=* 205) 200. The sample was followed by a checkerboard mask (100 ms) in the orientation task, while no mask was used in the number task.

**Figure 1.**
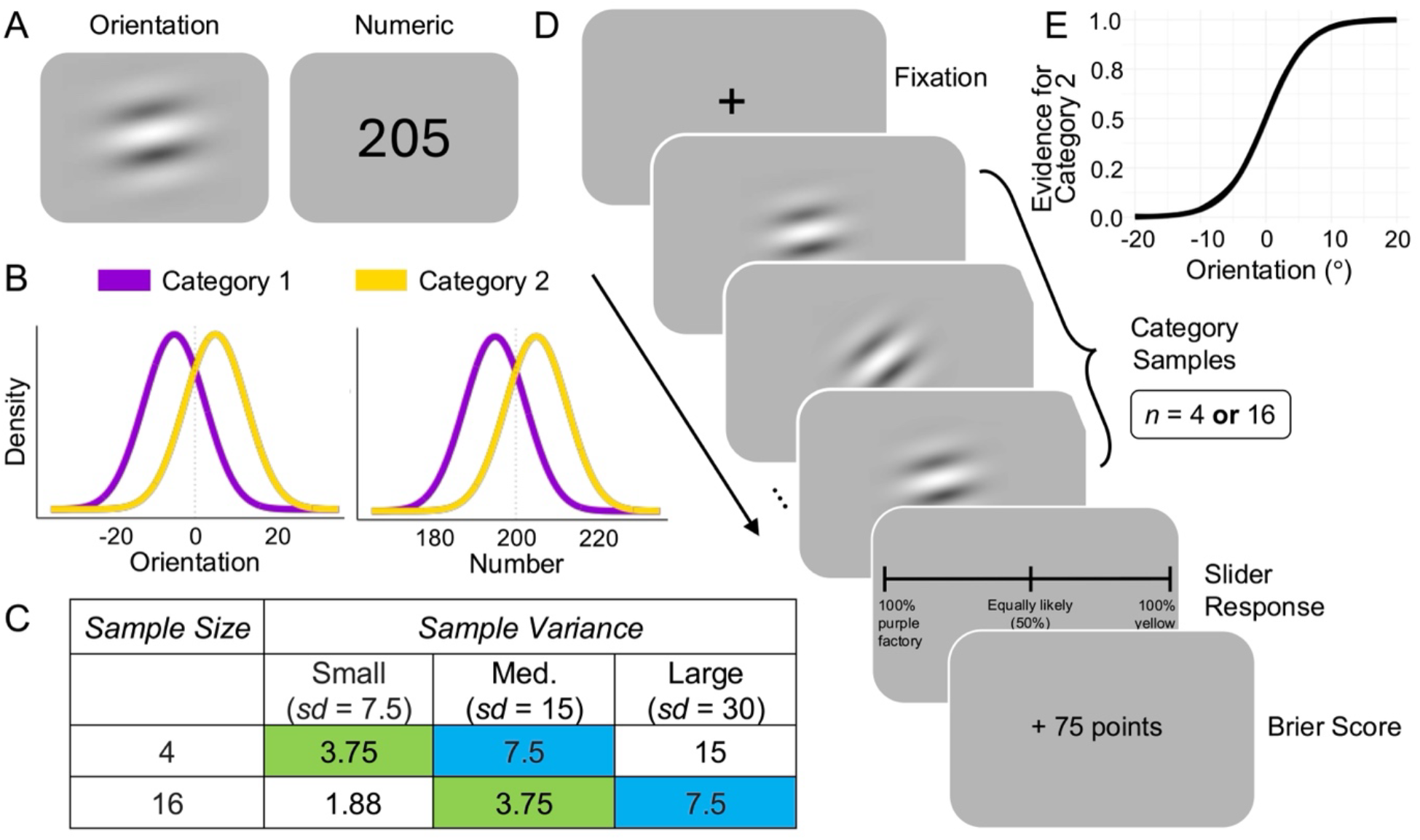
Experimental Procedure. *Note*. **(A)** Example orientation and number stimuli. **(B)** Example generating category distributions for Category 1 (purple) and Category 2 (yellow) in the small sample standard deviation condition. **(C)** Combinations of sample SD and sample size, where shaded cells of the same color represent conditions matched for standard error. **(D)** Example trial sequence (points displayed only during training). **(E)** Example changes in evidence for category 2 as a function of orientation stimulus values.

Participants were asked to judge the probability that the sequence had been generated by ‘category 1’ or ‘category 2’ on a slider scale. The categories were described as two factories labelled ‘yellow’ and ‘purple’. The slider had ticks at 0, 50 and 100 labelled ‘very confident purple factory’, ‘not sure’, ‘very confident yellow factory’ respectively (see ‘task instructions and cover story’ below for explanation of factory analogy). During testing, participants did not receive trial-by-trial feedback, but at the end of each block, they were informed of their total points earned out of a maximum possible score (e.g., “You scored X points out of 1200”).

Domain (orientation or number) was blocked and counterbalanced so that participants completed the categorization task with either orientation or number values first. Within each domain, we implemented a fully balanced experimental design in which we systematically varied the mean of the generating distribution (category 1 mean or category 2 mean), the standard deviation of the generating distributions (small, medium or large) and the number of observations (4 or 16 observations). This yielded 12 cells. We constructed number sequences for each cell of the design. To ensure that the presented sequences were representative of the respective generating distributions, sequences were generated to match the target SD exactly. Sample means were allowed to vary naturally but observations that fell ±3 SD from the underlying distribution mean were excluded. Participants completed 36 testing trials across three blocks in each domain. Each block included one trial from each of the 12 cells of the design, the order of which was randomized. **Figure 1C** summarizes the experimental design. After completing the first domain, participants completed training and testing in the second domain. The full experiment lasted approximately 30–40 minutes.

### Task Instructions and Cover Story

Participants received detailed instructions about the task structure and the nature of the generating distributions. A cover story was used to aid comprehension: participants were told that the “yellow” and “purple” factories produced numbers or patterns (oriented gratings) that differed on average. The standard deviation manipulation was described as the machines in the factories having different grades of reliability, where the lowest reliability machines produced highly variable output. Participants then saw an example sample with 10 observations from each mean and SD condition. They were also instructed on how to use the slider scale, with examples clarifying what different positions on the scale indicated (e.g. “I think there’s an 85% chance that the patterns belong to the purple factory” and “I think there’s a 60% chance that the patterns belong to the yellow factory”).

The scoring rule was then introduced, indicating that on each trial participants could earn up to 100 points. We told them that the number of points that they would earn would be determined by how far their selected probability was from the correct outcome. We used 3 examples to aid in the understanding of the scoring rule: “if you chose a point on the scale which is 80% towards the CORRECT factory, you will get 96 points”; “if you chose the middle of the scale, you will get 75 points”; “if you chose a point on the scale which is 80% towards the INCORRECT factory, you will get 36 points”. These examples were followed by: “this means that to get the most points, you should choose a point on the scale that accurately reflects how sure you are about an answer”. Participants were then told that at the end of the experiment, their points would be converted into a bonus payment (up to an additional £1.80). Below, we outline the underlying scoring rule in more detail (see ‘Incentivization’).

After the instructions, participants completed training with feedback (detailed below), after which they were introduced to the sample size manipulation. They were told that during the main experiment they would see sequences with either 4 or 16 observations, and that feedback would only be provided at the end of each block.

### Training

Participants completed four training blocks using 10-observation samples. The first three blocks used small, medium, and large SDs respectively, each consisting of four trials (two per generating category mean). The fourth block interleaved SDs randomly, with one trial per category mean. Feedback was provided on each training trial (“Correct” or “Incorrect” in green or red) along with the number of points earned.

### Incentivization

Participants earned points on each trial according to a Brier scoring rule, a strictly proper scoring rule that incentivizes jointly maximizing the accuracy of choice and confidence ratings. The Brier score for each trial was calculated as:

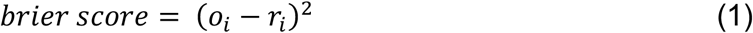

where *o*_*i*_ is the actual outcome (0 = category 1, 1 = category 2) and *r*_*i*_ is the participant’s probability judgment (scaled between 0 and 1) on trial *i*. Because lower Brier scores indicate better performance, participants’ points were computed as *points =* (1 − *brier score*) × 100. This scoring rule meant that participants could earn a maximum of 100 points per trial, with the highest rewards given for providing highly confident correct responses or low confidence incorrect responses. At the end of the experiment, total points were converted into a bonus payment at a rate of £0.02 per 100 points. The average bonus payment per participant was £1.12.

## Results

### Data cleaning

As outlined in our pre-registration (https://doi.org/10.17605/OSF.IO/Q538N), we excluded 28 participants who failed to discriminate between the generating categories better than chance, based on a binomial test of significance. We also pre-registered that we would remove trials where participants repeatedly chose the same point on the slider scale. However, due to concerns that this would create uneven numbers of trials across conditions, we post hoc decided not to exclude any trials.

Instead, we removed participants who selected the same point on the slider on more than 25% of consecutive trials. Using this criterion, we excluded an additional 10 participants. Because this exclusion criterion was not explicitly pre-registered, we confirmed that removing these 10 participants did not affect the main findings (see supplementary materials). Our final sample size was 85 participants, as recommended by our a priori power analysis.

### Data Analysis

We were interested in two main dependent variables: participants’ subjective probability judgments and the objective accuracy of these judgments. To facilitate interpretation of the probability responses, we transformed the raw probability values into a ‘confidence’ measure by calculating the distance of each point from the objective midpoint of the scale (i.e., 50%). This transformation represented perceived probabilities independently of the category decision (i.e., whether Category 1 or Category 2 was chosen) and ranged from 50 (chance) to 100 (100% confidence). The accuracy data, which was measured using the Brier score, ranged from 0 to 100.

We used linear mixed-effects models with sample size, sample standard deviation, and sample domain as fixed effects, and included a random intercept for each participant. We treated the independent variables as factors to test mean differences in the dependent measures across participants. Significant fixed effects are summarized below, and the full results are included in the supplementary materials. See **Table S2** for accuracy data and **Table S3** for confidence data.

### Accuracy Measure

We found a significant effect of sample size (*b* = 7.93, *t* = 4.5, *p* < .001) on the number of points participants received, where participants obtained more points for samples with a larger number of observations (*M*=83.3, *SD* =27.5) relative to samples with a smaller number of observations (*M*=76.0, *SD*=30.8). This is plausible given that larger sample size reduces uncertainty about the mean. We also found that the effect of small (*M*=87, *SD*=24.2), medium (*M*=79.4, *SD*=29), and large (*M*=72.4, *SD*=32.6) SD was significant. Compared to the small level, the medium level (*b* = −8.56, *t* = −4.84, *p* < .001) and large level (*b* = −14.14, *t* = −8, p < .001) were associated with fewer points. Again, this is plausible as greater variability increases uncertainty. We did not observe a significant effect of sample domain (*b* = −1.15, *t* = −0.65, *p* = .516) or any other significant interactions, indicating that accuracy in the number task was similar to the orientation task.

### Confidence Measure

We found a significant effect of sample size (*b* = 6.58, *t* = 7.16, *p* < .001) on our confidence measure, where participants were more confident about samples with a larger number of observations (*M*=84.2, *SD*=16.5) relative to samples with a smaller number of observations (*M*=79.9, *SD*=18.0). We also found that the effect of the small (*M*=85.1, *SD*=16.5), medium (*M*=81.3, *SD*=17.3), and large (*M*=79.8, *SD*=17.9) sample standard deviation was significant. Compared to the small level, the medium level (*b* = −1.92, *t* = −2.09, *p* = .037) and large level (*b* = −1.84, *t* = −2, p = .046) were associated with a decrease in confidence. These results are qualitatively similar to the accuracy measure reported above. In contrast to the accuracy measure, we also found a significant effect of sample domain (*b* = −2.33, *t* = −2.54, *p* = .011), where participants were more confident in number samples (*M*=83.1, *SD*=17.2) relative to orientation samples (*M*=81.0, *SD*=17.5). The interaction between sample size and standard deviation was significant for both the medium variability condition (*SD* = 15; *b* = –3.41, *t* = –2.63, *p* = .009) and the high variability condition (*SD* = 30; *b* = –5.41, *t* = –4.17, *p* < .001), indicating that the increase in confidence with sample size was stronger for less variable samples. In other words, the benefit of larger sample sizes for confidence judgments diminished as variability increased (see **Table S4** for estimated marginal means). This pattern suggests that when the evidence was less ‘reliable’ (i.e., higher variability), participants placed less weight on the number of observations when forming confidence judgments.

### Asymmetry in the Effect of Sample Size on Accuracy and Confidence

As described above, we observed an interesting asymmetry in the effect of sample size on accuracy and confidence. Participants were consistently more accurate with larger samples, and the increase in accuracy remained relatively stable across different levels of sample variability. In contrast, participants’ confidence increased with larger samples only when the sample variability was low or moderate. This meant that for high variance samples, accuracy was still higher for larger samples, but confidence did not differ. This dissociation suggests that confidence judgments may be less sensitive to total uncertainty when the evidence is highly variable, potentially reflecting limits in metacognitive sensitivity or integration under noisy conditions.

### The Effect of Standard Error

As shown in **Figure 2**, participants showed comparable accuracy and confidence across conditions matched for statistical uncertainty, defined as 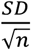 (i.e. standard error). To quantify this pattern, we conducted a series of planned pairwise comparisons. We found that for conditions with a low standard error (3.75), there was no difference in accuracy for either the number (*estimated difference* =1.04, *z-ratio* = 0.59, *p* = 0.557) or orientation samples (*estimated difference* = −0.63, *z-ratio* = −0.36, *p* = 0.723). Similarly, for conditions with a high standard error (7.5), there was no difference in accuracy for either the number (*estimated difference* = 2.22, *z-ratio* = 1.26, *p* = 0.209) or orientation (*estimated difference* = −1.78, *z-ratio* = −1.01, *p* = 0.314) samples.

**Figure 2.**
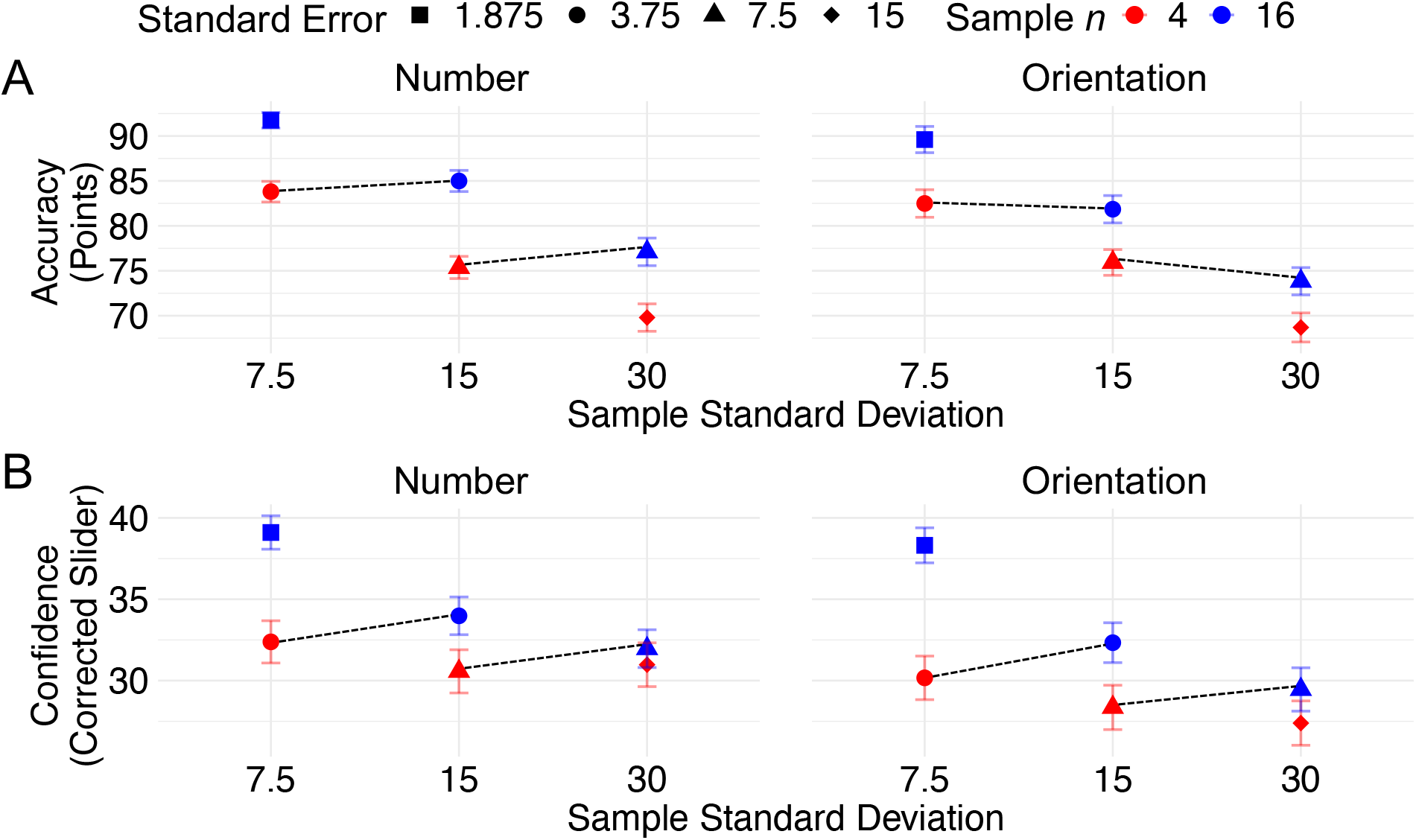
Accuracy and Confidence Across Conditions. *Note*. **(A)** Accuracy for number (left) and orientation (right) tasks as a function of sample size (colors) and sample standard deviation (x-axis). **(B)** As in (A), but for confidence.

In terms of confidence, for the low standard error condition, we found a significant difference for the orientation samples, where samples with lower variance and fewer observations were associated with higher confidence (*estimated difference* = 2.27, *z-ratio* = 2.48, *p* = 0.013). No significant difference in confidence was found for the number of samples in the low standard error condition (*estimated difference* = 1.25, *z-ratio* = 1.36, *p* = 0.174). For the high standard error condition, there were no significant differences in confidence for either the number (*estimated difference* = 1.25, *z-ratio* = 1.36, *p* = 0.174) or orientation samples (*estimated difference* = 1.02, *z-ratio* = 1.11, *p* = 0.267).

### Model Specification and Fitting

We compared seven models that differed in their assumptions about how participants used sample statistics to form their choice and confidence judgments. The models differed in the sophistication of the cognitive computations assumed to be underlying responses, ranging from simple reliance on the sample mean to more optimal strategies that approximated the computation of a posterior probability. All models assumed that participants responses were influenced by the sample mean, such that lower sample means were associated with higher-confidence judgments for category 1, and higher sample means were associated with higher-confidence judgments for category 2. The models differed in how this relationship was modulated by uncertainty-related factors such as sample size, variability, or standard error.

Each model was fit separately to the orientation and number data, allowing us to examine whether participants used similar computational strategies across domains. To capture the full range of slider responses (0 to 1), we modelled responses using a zero-one inflated beta distribution. This distribution includes a point mass at 0, a point mass at 1, and a continuous beta-distributed component (*μ*_*i*_) for intermediate values. The same trial-level predictors that influenced the mean of the beta distribution were also used to model the probability of extreme responses (0 or 1), allowing us to account for how sample characteristics affected the entire response distribution. For each model, the mean of the beta-distributed component for trial *i*, denoted *μ*_*i*_, was modelled as follows:

I. Mean model:

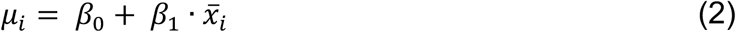

where 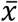 was the sample mean.
II. Sample size model:

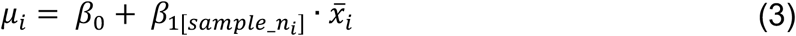

This model allowed the effect of the sample mean to differ depending on the number of observations in the sample, *sample_n*.
III. Standard deviation model:

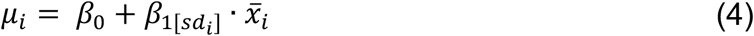

This model allowed the effect of the sample mean to differ depending on the standard deviation of the sample, *sd*.
IV. Additive model:

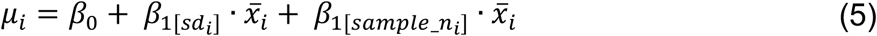

This model allowed both sample standard deviation and sample size to independently modulate the effect of sample mean, assuming additive (non-interacting) contributions of each factor.
V. Interactive model:

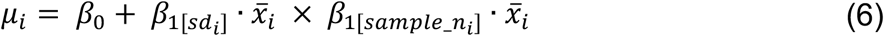

This model allowed the effect of the sample mean to depend jointly on both sample size and standard deviation. This allowed the model to capture dependencies between the two sources of uncertainty, for example whether the influence of sample size on confidence is stronger when variability is low.
VI. Standard error model:

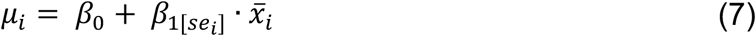

This model allowed the effect of the sample mean to differ depending on the standard error of the sample, 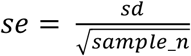., which captures statistical uncertainty in the sample mean. It thus accounts for the fact that the same sample mean may be more or less informative depending on the precision of the sample. Although this model incorporates key statistical features of the sample, it is not Bayesian because it does not assume prior knowledge of the generating category distributions, nor does it involve computing likelihoods or a posterior probability.
VII. Bayesian model:

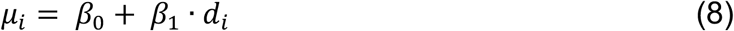

where *d* is the posterior probability ratio which was calculated as:

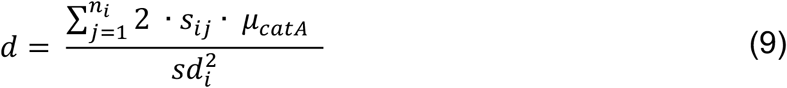

where *s*_*ij*_ represented the value of the *j*^th^ observation in the sample, *n* was the number of observations in the sample, *μ*_*catA*_ was the mean of the generating distribution for Category 1 and *sd* was the sample standard deviation (constrained to match the true standard deviation of the generating distribution) on trial *i*. The resulting value of *d* can be interpreted as a measure of how much more likely it is that the sample came from Category 1, such that the higher the value of *d*, the more evidence in favor of Category 1. This model approximated an ideal observer strategy under Bayesian inference.

All models were fit using the brms package in R, with four chains, 4000 total iterations per chain (1000 warmup), and weakly informative priors. For model comparison, we computed the expected log pointwise predictive density using leave one-out cross validation (*elpd_loo*) via Pareto-smoothed importance sampling for each model. This method estimates each model’s expected predictive accuracy by assessing how well it predicts each observation when that observation is left out of model fitting. The standard error for *elpd_loo* was computed as the square root of the sum of the variances of the pointwise *elpd_loo* values across observations. This reflects the uncertainty in the model’s out-of-sample predictive accuracy. To further facilitate model comparison, we also computed Akaike weights, which normalize *elpd_loo* values across the full model set to reflect each model’s relative predictive performance. Model weights incorporate both the magnitude and uncertainty of the *elpd_loo* estimates across models to provide a normalized probability that each model is the best among the set. As a result, a model may have a slightly better (lower) LOO score but still receive a lower weight if the difference is small relative to the estimated variability across models.

To visualize model predictions, we generated a simulated dataset by drawing a sample from the posterior distribution of parameters and then simulating a new response from the likelihood function. This posterior-predictive approach incorporates both parameter uncertainty and observation-level noise, and represents a single plausible set of participant responses under the fitted model. For model predictions that include a credible interval, we instead computed 95% credible intervals across posterior draws for each observation using the expected value of the response. These intervals reflect uncertainty in the model’s predicted mean response due to parameter uncertainty, but do not include observation-level noise, as no responses are drawn from the likelihood in this case.

### Model Comparison Results

The mean-only model (model i; Equation 2) assumed that participants relied solely on the sample mean to make their confidence judgments, without adjusting for uncertainty in that mean. Table 2 shows that this model performed the worst across both the orientation and number tasks. It predicted the steepest changes in confidence in high-uncertainty conditions, where sample means tended to be more extreme due to greater variability. As a result, this model systematically overpredicted confidence for high-uncertainty samples and underpredicted confidence for low-uncertainty samples, as shown in **Supplementary Figure 2**. The poor performance of this model suggests that participants did not rely on a simple heuristic based solely on the sample mean, but instead adjusted their responses according to the uncertainty associated with using the sample mean to estimate the population mean.

**Table 2.**
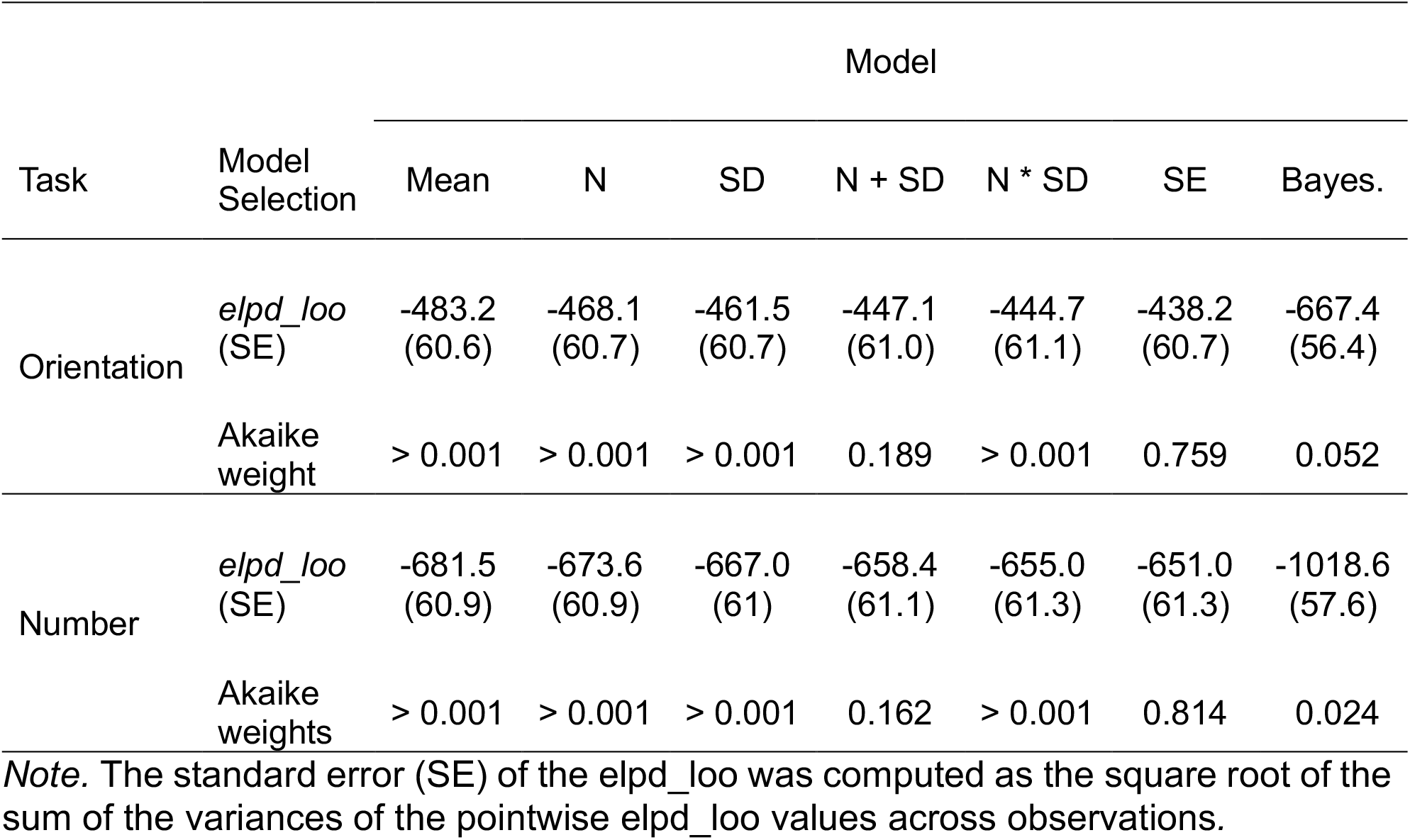
Model Comparison.

| Task | Model Selection | Model |  |  |  |  |  |  |
| --- | --- | --- | --- | --- | --- | --- | --- | --- |
|  |  | Mean | N | SD | N + SD | N * SD | SE | Bayes. |
| Orientation | *elpd_loo* (SE) | -483.2 (60.6) | -468.1 (60.7) | -461.5 (60.7) | -447.1 (61.0) | -444.7 (61.1) | -438.2 (60.7) | -667.4 (56.4) |
|  | Akaike weight | > 0.001 | > 0.001 | > 0.001 | 0.189 | > 0.001 | 0.759 | 0.052 |
| Number | *elpd_loo* (SE) | -681.5 (60.9) | -673.6 (60.9) | -667.0 (61) | -658.4 (61.1) | -655.0 (61.3) | -651.0 (61.3) | -1018.6 (57.6) |
|  | Akaike weights | > 0.001 | > 0.001 | > 0.001 | 0.162 | > 0.001 | 0.814 | 0.024 |

*Note.* The standard error (SE) of the *elpd\_loo* was computed as the square root of the sum of the variances of the pointwise *elpd\_loo* values across observations.

Models that included only one aspect of uncertainty in the sample mean, either sample size (model ii; Equation 3) or sample standard deviation (model iii; Equation 4), provided a modest improvement over the mean-only model. As shown in **Figure 2**, responses depended both on the number of observations and the variability of the sample. Because the sample size (model ii) and sample standard deviation (model iii) models each account for only a single source of uncertainty, they were unable to reproduce this joint influence. For example, **Supplementary Figure 3** shows that the sample size model (ii) did not capture changes in confidence as a function of sample variability and **Supplementary Figure 4** shows that the standard deviation model (iii) did not capture the increase in confidence for samples with a larger number of observations.

**Table 2** further shows that models which included both sample size and variability, either as additive (model iv; Equation 5) or interactive predictors (model v; Equation 6), showed further improvements in model fit. These models reproduced the observed pattern of steep changes in confidence with sample means for large, low-variability samples, and shallower changes for small, high-variability samples. This behavior is visible in **Supplementary Figures 5 and 6**. However, despite their flexibility, these models treated sample size and variability as separate inputs rather than integrating them into a unified estimate of uncertainty. As such, they provided a useful approximation but lacked the parsimony of the standard error model (described below).

As reported in **Table 2**, the best-fitting model included sample standard error as a predictor that modulated the effect of the sample mean (model vii; Equation 8). In addition to being the best-fitting model, **Figure 5** shows that the model predictions provided a close fit to the data, capturing changes in confidence and accuracy across different levels of sample variability, sample size, and task domain. By scaling the influence of the sample mean according to the standard error, which combined both sample size and variability, this model captured the interaction between sample mean and sample uncertainty. The left panels of **Figure 5** demonstrate that the model successfully reproduced the steeper relationship between sample mean and confidence for samples with a small standard error. In contrast, the right panels of **Figure 5** demonstrate that the model also successfully reproduced the flatter relationship between sample mean and confidence for samples with a large standard error. The success of the standard error model in capturing behavior suggests that participants did not treat sample size and variance as separate or arbitrarily interacting cues but instead integrated them in a manner that approximated the normative standard error of the mean.

**Figure 4.**
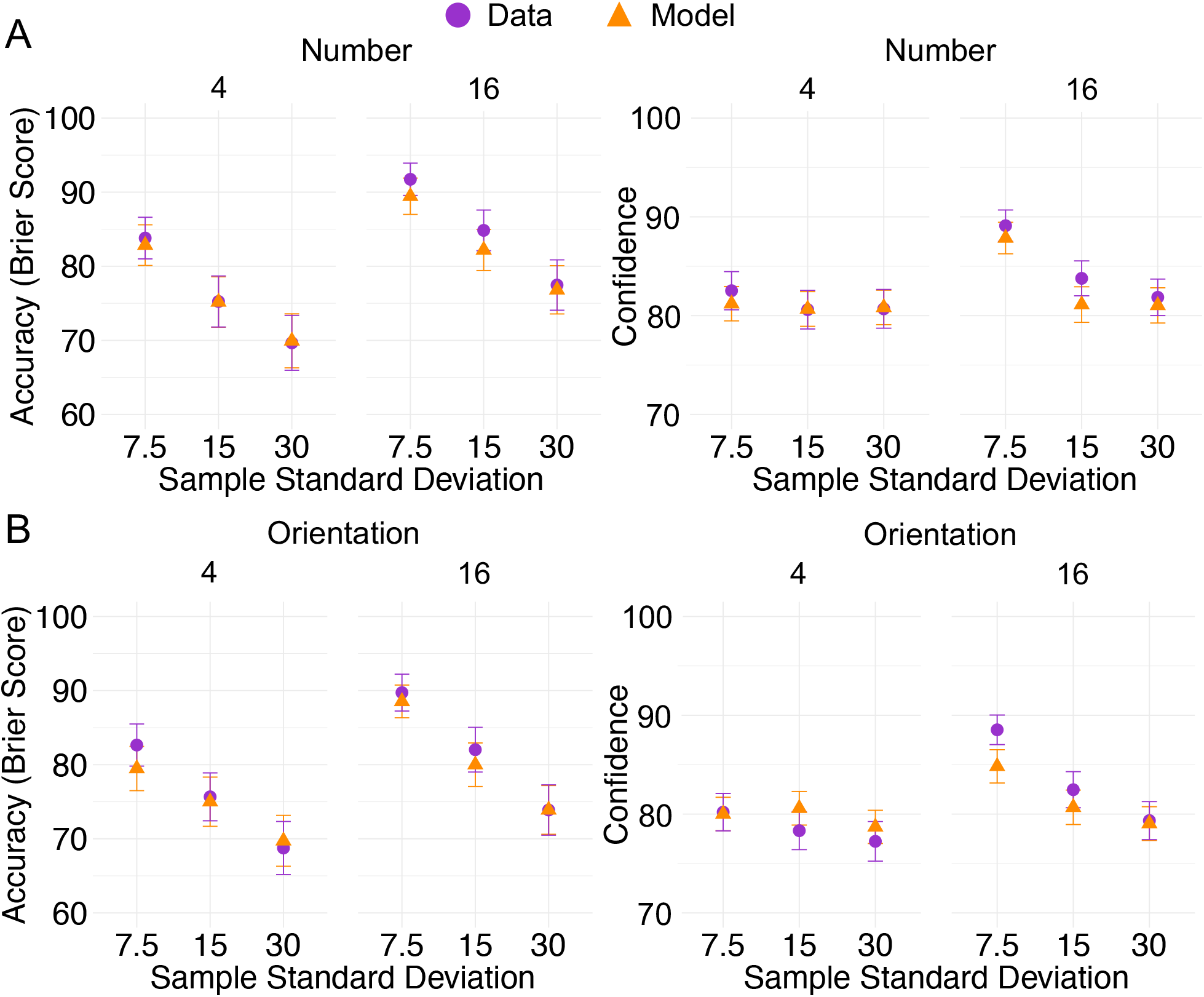
Fit of Standard Error Model to Accuracy and Confidence Data. *Note*. Model predictions from the standard error model (orange) and data (purple) for the orientation **(A)** and number **(B)** task. Error bars reflect ±1 SEM across participants within each condition. For the model, we generated a simulated dataset from the posterior predictive distribution (i.e., draws from the joint posterior over parameters and observations).

**Figure 5.**
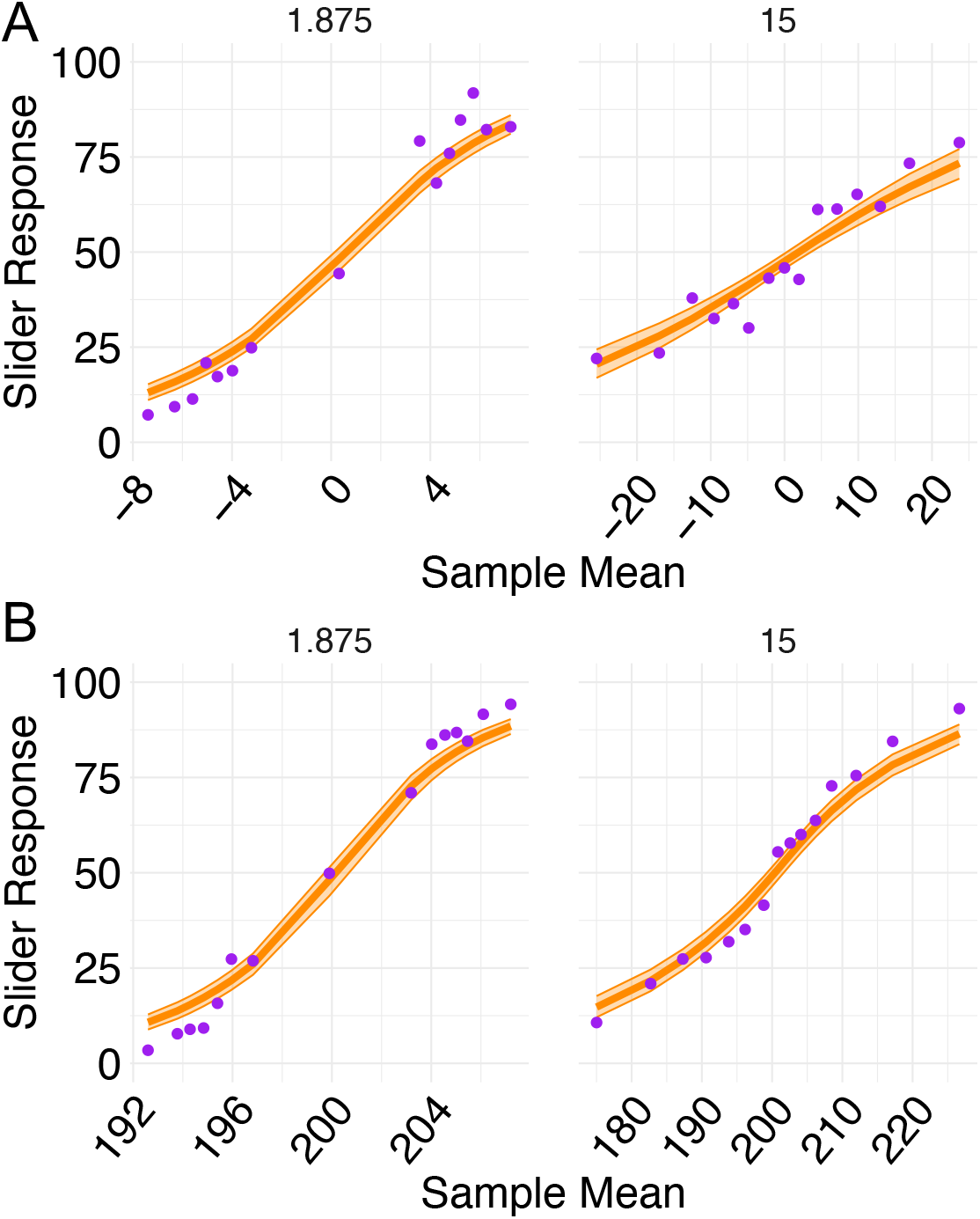
Standard Error Model Across Sample Means. *Note*. Model predictions (orange) and empirical data (purple) for the standard error model, shown separately for the orientation task **(A)** and the number task **(B)**. Confidence increases with the sample mean, and the slope of this relationship varies systematically with the standard error: steeper when uncertainty is low, flatter when uncertainty is high. Error bars indicate 95% credible intervals for the model’s expected values, computed from posterior samples.

The Bayesian model (model vi; Equation 7) assumed participants computed the posterior probability of category membership by integrating all available sample statistics (sample mean, sample variance, and sample size) with prior knowledge of the category distributions. As shown in **Supplementary Figure 7**, this model provided a reasonable fit but it failed to fully capture how confidence varied across levels of uncertainty. Specifically, the Bayesian model predicted more gradual changes in confidence in high-uncertainty (high standard error) samples than were observed in the empirical data. Participants were often more confident, and more often correct, than the model anticipated for these samples. This mismatch was especially evident in the Brier score, which penalizes underconfident responses to correct answers. As a result, even though the Bayesian model captured some general patterns in the confidence data, it significantly underestimated participants’ accuracy for high-uncertainty samples.

### Domain-General Model

To assess whether participants relied on similar strategies across the orientation and number tasks, we used the best-performing model, the standard error model, for additional model comparison. As shown in **Supplementary Figure 9**, we fit this model in two ways: first, as a domain-general model, jointly to both datasets under the assumption of shared parameters; and second, as domain-specific models, estimated separately for each task (as described above). Model comparison was again conducted using the LOO criterion. The difference in expected *elpd_loo* between the domain-specific and domain-general models was small (*elpd_diff* = –42.02, *SE* = 122.29). The corresponding z-score (*z* = –0.34) indicated no reliable evidence that the domain-specific models provided a better fit than the domain-general model. This result suggests that a single parameterization of the standard error model may be sufficient to account for participants’ behavior across both tasks.

## Discussion

The subjective sense of certainty that people have in their decisions, referred to as decision *confidence*, plays a central role in guiding behavior. It supports adaptive control by influencing whether individuals seek additional information (Desender et al., 2018; van den Berg, Zylberberg, et al., 2016), how they prioritize competing task (Aguilar-Lleyda et al., 2020), or allocate cognitive resources (Boldt et al., 2019). These functions are most effective when confidence is well calibrated, meaning that subjective certainty aligns with the objective uncertainty present in the evidence on which the decision is based. In this study, we examined how choice and confidence judgments reflect two key sources of statistical uncertainty: the number of observations in a sample (sample size) and the variability of those observations (sample variance). Importantly, we tested this in two distinct domains, using visual orientation stimuli and number stimuli to probe the generalizability of these computations.

### Evidence for Statistically Grounded Confidence Computations

Our results showed that participants’ accuracy and confidence were influenced by both sample size and sample variance across domains. Overall, participants were more accurate and more confident in samples that were less variable and had a larger number of observations, consistent with prior findings that confidence reflects both the strength and reliability of evidence (Boldt et al., 2017; Spence et al., 2016). Participants also appeared to account for the trade-off between these factors, showing comparable levels of confidence and accuracy in conditions where greater variability was offset by a larger number of observations. Specifically, when samples were matched in terms of statistical uncertainty (i.e., had the same standard error of the sample mean), there were no differences in accuracy and only very small differences in confidence, only in the orientation task. This finding suggests that confidence aligns with objective statistical uncertainty, and provided some preliminary evidence that people use statistically guided strategies to compute their confidence.

To more directly test this hypothesis, we used computational modeling to compare how well different models explained participants’ confidence judgments. A model that relied solely on the sample mean performed poorly, indicating that participants did not treat all sample means equally but adjusted their confidence according to how uncertain those means were. Models that treated sample size and sample variance as independent, additive, or even interacting factors were outperformed by a model that modulated the influence of the sample mean based on the sample’s standard error. These modeling results reinforce the behavioral trends and further suggest that people use statistically guided strategies to compute their confidence, rather than relying on heuristic processes that over- or underweight individual aspects of uncertainty, or that fail to account for the trade-off between sample size and variance.

Although participants’ judgments reflected sensitivity to statistical uncertainty, such as the standard error, we did not find evidence that they engaged in full Bayesian inference. The Bayesian model that incorporated prior knowledge of the generating category distributions and computed a posterior probability ratio using the full set of sample statistics did not outperform the standard error model. While this contrasts with studies suggesting that confidence sometimes aligns with Bayesian inference (e.g., Fleming & Daw, 2017; Meyniel et al., 2015), it is consistent with research showing that people use statistical cues without fully implementing Bayesian computations (Navajas et al., 2017; Sanders et al., 2016). Our findings extend this literature by demonstrating that even when both sample size and variance are manipulated, participants’ behavior aligns more closely with approximate statistical strategies than with fully optimal Bayesian inference.

One reason for the Bayesian model’s poor performance may be the complexity of the optimal Bayesian decision rule in our task. Unlike other tasks, which involve a single prior distribution over the population mean (as in Olschewski & Scheibehenne, 2024), our categorization task involved two discrete categories, each defined by its own prior distribution. Making an optimal decision in this context required computing the likelihood of the observed sample under each category, then comparing these to compute a posterior probability ratio. This type of decision rule may involve computations that are more cognitively complex. In this context, participants may have defaulted to a more tractable strategy, approximating uncertainty using the standard error of the sample mean. As such, the standard error model may have served as a computationally efficient ‘good-enough’ solution that closely tracked the normative trade-offs without requiring full probabilistic inference.

### Limitations of our Modeling Assumptions

Importantly, the poor performance of the Bayesian model should not be taken as evidence against Bayesian accounts more broadly. The specific implementation we tested involved several simplifying assumptions, most notably, that participants had perfect knowledge of the generating category distributions, and that both their internal representations of the sample statistics and the computational process itself were noise-free. In practice, participants may have used approximate or biased versions of the category means, or formed noisy internal representations of the sample statistics, noise which they may have explicitly taken into account during decision-making. By fixing these parameters, our model was designed to isolate the core structure of Bayesian inference, without additional complexity. Future work could explore more flexible Bayesian frameworks that estimate subjective priors and incorporate internal noise, to better capture individual variability and potential departures from optimality.

A more general limitation of our modeling approach is that we assumed participants computed the true sample mean from the full set of observations on each trial. This assumption may not fully reflect participants’ actual strategies, as there is evidence that people often rely on simplified or biased estimators, such as running averages or weighting recent items more heavily due to recency effects, or initial items due to primacy biases. Incorporating such mechanisms into the generative model would undoubtedly offer a richer account of how evidence is integrated over time. However, our primary aim was not to model the precise process by which participants computed summary statistics, but rather to compare different computational frameworks—Bayesian, frequentist, and heuristic—for how such summaries are used to form confidence judgments. Including alternative encoding models for the sample mean would have vastly expanded the model space and made direct comparison across computational strategies more difficult. We therefore opted to hold the sample summary constant and focus on characterizing the inferential computations that operate on that input.

### Dissociation Between Confidence and Accuracy

One particularly interesting finding was the systematic dissociation between confidence and accuracy, where accuracy was more sensitive to sample variance than confidence. While we observed no interaction between sample size and variability in participants’ accuracy, such an interaction emerged clearly in the confidence data. Specifically, participants’ confidence increased with sample size more strongly when variability was low than when it was high. This suggests that people are less inclined to scale confidence with evidence quantity when the evidence itself is noisy, even when doing so leads to more accurate decisions. This dissociation highlights that confidence reflects perceived evidence reliability (Boldt et al., 2017; Spence et al., 2016), which may diverge from objective accuracy. This has important implications for models of confidence. It illustrates that confidence is not merely a readout of accuracy or performance, but instead arises from a different evaluation process. Future models should accommodate the idea that confidence and accuracy, while often aligned, may be influenced by different dimensions of uncertainty depending on the task conditions.

### Uncertainty across Domains

Overall, our findings align broadly with the existence of a domain-general system for uncertainty computation. Across both the orientation and number tasks, the same pattern of effects emerged: confidence tracked statistical uncertainty, and the same computational model provided the best fit to behavior in both domains. Further modeling work confirmed that the same parameterization could account for responses across both tasks. This cross-domain consistency suggests that confidence may rely on shared computational principles that generalize across task types.

Prior research on the question of generalization across domains has produced mixed findings. Some studies support domain-general mechanisms (Ais et al., 2016; Faivre et al., 2018; Fleck et al., 2006; Heereman et al., 2015; Mazancieux et al., 2020; Morales et al., 2018; Pleskac & Busemeyer, 2010; Rouault et al., 2018; Song et al., 2011; West et al., 2023) while others indicate domain-specific metacognitive abilities (Baird et al., 2013, 2015; Faivre et al., 2018; Fitzgerald et al., 2017; Garfinkel et al., 2016; Kelemen et al., 2000; McCurdy et al., 2013; Pannu et al., 2005; Schnyer et al., 2004; Valk et al., 2016). Our modeling approach offers a complementary perspective: Unlike approaches that focus solely on observed confidence ratings or neural patterns, it provides a mechanistic lens through which to assess whether confidence across tasks is supported by shared computational processes. The clear behavioral and computational convergence observed here suggests that across perceptual and cognitive domains, confidence judgments may be governed by a common underlying architecture.

### Contrast with Prior Work

Finally, our findings contrast with the results of Olschewski and Scheibehenne (2024). In their experiment, participants estimated the mean of a sequence of numbers (similar to our number condition). They found that participants’ mean estimates were less accurate in high-variance, large-sample-size conditions that were matched in standard error to low-variance, small-sample-size conditions. They attributed this discrepancy to heightened epistemic uncertainty, proposing that cognitive imprecision increases when participants must integrate a larger number of highly variable number values, making it harder to form and maintain an accurate summary representation.

One plausible explanation for the divergent findings between our results and those of Olschewski and Scheibehenne (2024) lies in differences in task demands and response format. In their study, participants estimated the mean of a sequence of numbers and placed a monetary bet on the accuracy of their response. This betting component may have introduced additional risk-related considerations, amplifying perceived uncertainty and promoting more cautious or risk-averse responses under high variability. In contrast, our task required categorical judgments (i.e., selecting category 1 or 2) and explicit probability estimates, potentially encouraging more direct, statistically grounded confidence assessments. These differences in the task and the answer format might explain why we observed more consistent and calibrated confidence in large sample size, high-variance samples, and why our participants’ behavior tracked statistical uncertainty more closely. More broadly, these conflicting results show how task structure, including the number of prior distributions and the nature of the response, can shape the computational strategies people use to evaluate uncertainty.

## Conclusion

Together, these findings offer new insights into how people compute confidence from uncertain evidence. By orthogonally manipulating sample size and variability across two tasks, we found that participants used a strategy that approximated key statistical principles. Confidence closely tracked the standard error of the sample mean, reflecting both evidence quantity and variability, consistent with a structured but non-Bayesian approach to uncertainty. Importantly, this strategy appeared to generalize across the number and orientation task, supporting the idea of a domain-general mechanism for confidence computation. Our results suggest that confidence is not simply a readout of accuracy, but a distinct and nuanced assessment of evidence uncertainty, one that may rely on shared, computationally tractable principles across tasks and contexts. Future work should build on these findings to further characterize the limits of this generalizability and the neural mechanisms that support flexible uncertainty tracking.

## Supporting information

All supplemental

## Notes

### Competing Interest Statement

The authors have declared no competing interest.

### Funding

- Helmholtz Information and Data Science Academy

