## Supplementary material for "Confidence Judgments Reflect the Standard Error of Noisy Evidence Samples Across Domains": All supplemental

### Supplementary Materials

#### Checking Behavioral Results Are Robust to Exclusion Decisions

Because we did not preregister the decision to exclude participants who selected the same point on the slider scale on more than 25% of consecutive trials, we verified that this exclusion did not affect the main behavioral trends. In the following sections, we report key behavioral results using data from all 95 participants, showing that the findings are consistent with those based on the reduced sample.

##### *Accuracy Measure*

We found a significant effect of sample size ($b$ = 8.73, $t$ = 5.04, $p$ < .001) on the number of points participants received, where participants obtained more points for samples with a larger number of observations ($M$=83.2 $SD$ =28.3) relative to samples with a smaller number of observations ($M$=75.7, $SD$=32). This is plausible given that larger sample size reduces uncertainty about the mean. We also found that the effect of small ($M$=87.1, $SD$=24.8), medium ($M$=79.3, $SD$=30), and large ($M$=71.9, $SD$=33.8) SD was significant. Compared to the small level, the medium level ($b$ = -8.02, $t$ = -4.64, $p$ < .001) and large level ($b$ = -13.72, $t$ = -7.92, $p$ < .001) were associated with fewer points. Again, this is plausible as greater variability increases uncertainty. We did not observe a significant effect of sample domain ($b$ = 0.61, $t$ = 0.36, $p$ = .723) or any other significant interactions.

##### *Confidence Measure*

We found a significant effect of sample size ($b$ = 5.72, $t$ = 6.65, $p$ < .001) on our confidence measure, where participants were more confident about samples with a larger number of observations ($M$=85.1, $SD$=16.6) relative to samples with a smaller number of observations ($M$=81.3, $SD$=18.2). The result showed that the effect of small ($M$=86.1, $SD$=16.5), medium ($M$=82.6, $SD$=17.5), and large ($M$=81, $SD$=18.2) standard deviation was significant. Compared to the small level, the medium level ($b$ = -1.94, $t$ = -2.26, $p$ = .024) and large level ($b$ = -2.11, $t$ = -2.46, $p$ = .014) were associated with a decrease in confidence. We found a significant effect of sample domain ($b$ = -2.57, $t$ = -2.99, $p$ = .003), where participants were more confident for number samples ($M$=84.3, $SD$=17.2) relative to orientation samples ($M$=82.1, $SD$=17.8). The interaction between sample size and standard deviation was significant for both the medium variability condition ($SD$ = 15; $b$ = –2.99, $t$ =-2.46, $p$ = .014) and the high variability condition ($SD$ = 30; $b$ = –4.70, $t$ = -3.86, $p$ < .001), indicating that the increase in confidence with sample size was stronger for less variable samples (see **Table S1**).

**Table S1**

*Bonferroni-Adjusted Pairwise Comparisons of Sample Size by SD and Modality (95 Participants)*

| | Accuracy | | |
|---|---|---|---|
| | 7.5 | 15 | 30 |
| Orientation | 6.91 ** | 8.06 ** | 4.27 ** |
| Number | 8.73 ** | 9.86 ** | 6.86 ** |
| | Confidence | | |
| | 7.5 | 15 | 30 |
| Orientation | 7.66 ** | 3.71 *** | 1.69 * |
| Number | 5.72 ** | 2.73 ** | 1.02 |

*Note.* Reported marginal means represent the contrast between sample size 16 and sample size 4. p-values have been adjusted using the Bonferroni correction to account for multiple comparisons. *$p$ < 0.050, **$p$ < 0.010, ***$p$ < 0.001

#### The Effect of Standard Error

Participants showed comparable accuracy and confidence across conditions matched for statistical uncertainty, defined as $\frac{SD}{\sqrt{n}}$ (i.e., standard error). To quantify this pattern, we conducted a series of planned pairwise comparisons. We found that for conditions with a low standard error (3.75), we found no difference in accuracy for either the number (*estimate* =1.84, *z-ratio* = 1.06, *p* = 0.288) or orientation samples (*estimate* = -0.52, *z-ratio* = -0.3, *p* = 0.765). Similarly, for conditions with a high standard error (7.5), there was no difference in accuracy for either the number (*estimate* = 1.17, *z-ratio* = 0.68, *p* = 0.498) or orientation (*estimate* = -1.47, *z-ratio* = -0.85, *p* = 0.396) samples.

In terms of confidence, for the low standard error condition, we found a significant difference for the orientation samples, where samples with less noise and fewer observations were associated with more confidence (*estimate* = 1.92, *z-ratio* = 2.23, *p* = 0.025). No significant difference in confidence was found for the number samples in the low standard error condition (*estimate* = 0.79, *z-ratio* = 0.92, *p* = 0.360). For the high standard error condition, there were no significant differences in confidence for either the number (*estimate* = 0.85, *z-ratio* = 0.99, $p$ = 0.323) or orientation samples (*estimate* = 0.58, *z-ratio* = 0.67, $p$ = 0.501).

#### Results From Full Linear Mixed Effect Models for Accuracy and Confidence

**Table S2** and **Table S3** report the full results from the linear mixed-effects models predicting accuracy (Brier scores) and confidence (corrected slider responses), respectively.

**Table S2**

*Mixed-Effects Model Results for Accuracy Data*

| Predictors | Estimates | *CI* | *p* |
|---|---|---|---|
| (Intercept) | 83.8 | 81.18 – 86.42 | **<0.001** |
| sample n [16] | 7.93 | 4.47 – 11.40 | **<0.001** |
| sd [15] | -8.56 | -12.02 – -5.09 | **<0.001** |
| sd [30] | -14.14 | -17.60 – -10.67 | **<0.001** |
| modality [visual] | -1.15 | -4.61 – 2.32 | 0.516 |
| sample n [16] × as sd15 | 1.67 | -3.23 – 6.57 | 0.505 |
| sample n [16] × as sd30 | -0.13 | -5.03 – 4.77 | 0.957 |
| sample n [16] × modality [visual] | -0.85 | -5.76 – 4.05 | 0.732 |
| sd [15] × modality [visual] | 1.57 | -3.33 – 6.47 | 0.529 |
| sd [30] × modality [visual] | 0.24 | -4.66 – 5.14 | 0.924 |
| (sample n [16] × as sd15) × modality [visual] | -2.38 | -9.32 – 4.55 | 0.5 |
| (sample n [16] × as sd30) × modality [visual] | -1.81 | -8.74 – 5.12 | 0.608 |
| Random Effects | | | |
| $\sigma^2$ | 797.11 | | |
| $\tau_{00\ ID}$ | 18.91 | | |
| ICC | 0.02 | | |
| $N_{ID}$ | 85 | | |
| Observations | 6120 | | |
| Marginal $R^2$ / Conditional $R^2$ | 0.058 / 0.079 | | |

*Note.* CI values refer to 95% confidence intervals calculated using the profile likelihood method. Significance values were obtained using the Satterthwaite approximation to calculate the degrees of freedom for the t-distribution based on the estimated variance-covariance matrix of the model parameters (Lüdecke, 2022).

**Table S3**

*Mixed-Effects Model Results for Confidence Data*

| Predictors | Estimates | *CI* | *p* |
|---|---|---|---|
| (Intercept) | 32.53 | 30.29 – 34.76 | **<0.001** |
| sample n [16] | 6.58 | 4.78 – 8.38 | **<0.001** |
| sd [15] | -1.92 | -3.72 – -0.12 | **0.037** |
| sd [30] | -1.84 | -3.64 – -0.04 | **0.046** |
| modality [visual] | -2.33 | -4.13 – -0.53 | **0.011** |
| sample n [16] × as sd15 | -3.41 | -5.96 – -0.87 | **0.009** |
| sample n [16] × as sd30 | -5.41 | -7.96 – -2.87 | **<0.001** |
| sample n [16] × modality [visual] | 1.75 | -0.79 – 4.30 | 0.177 |
| sd [15] × modality [visual] | 0.04 | -2.50 – 2.59 | 0.974 |
| sd [30] × modality [visual] | -1.12 | -3.67 – 1.43 | 0.389 |
| (sample n [16] × as sd15) × modality [visual] | -0.77 | -4.37 – 2.83 | 0.673 |
| (sample n [16] × as sd30) × modality [visual] | -0.82 | -4.42 – 2.78 | 0.655 |
| Random Effects | | | |
| $\sigma^2$ | 215 | | |
| $\tau_{00\ ID}$ | 74.97 | | |
| ICC | 0.26 | | |
| $N_{ID}$ | 85 | | |
| Observations | 6120 | | |
| Marginal $R^2$ / Conditional $R^2$ | 0.041 / 0.289 | | |

*Note.* CI values refer to 95% confidence intervals calculated using the profile likelihood method. Significance values were obtained using the Satterthwaite approximation to calculate the degrees of freedom for the t-distribution based on the estimated variance-covariance matrix of the model parameters (Lüdecke, 2022).

#### Pairwise Comparisons of Sample Size

**Table S4** shows Bonferroni-corrected pairwise comparisons of sample size levels, computed separately for each SD and modality combination using estimated marginal means.

**Table S4**

*Bonferroni-Adjusted Pairwise Comparisons of Sample Size by SD and Modality (85 Participants)*

| | Accuracy | | |
|---|---|---|---|
| | 7.5 | 15 | 30 |
| Orientation | 7.08 ** | 6.36 ** | 5.13 ** |
| Number | 7.93 ** | 9.60 ** | 7.80 ** |
| | Confidence | | |
| | 7.5 | 15 | 30 |
| Orientation | 8.33 ** | 4.15 ** | 2.10 * |
| Number | 6.58 ** | 3.17 ** | 1.16 |

*Note.* Reported marginal means represent the contrast between sample size 16 and sample size 4. p values have been adjusted using the Bonferroni correction to account for multiple comparisons. **p* < 0.050, ***p* < 0.010, ****p* < 0.001

**Figure S1**

*Histogram of Confidence Responses across SD and N conditions*

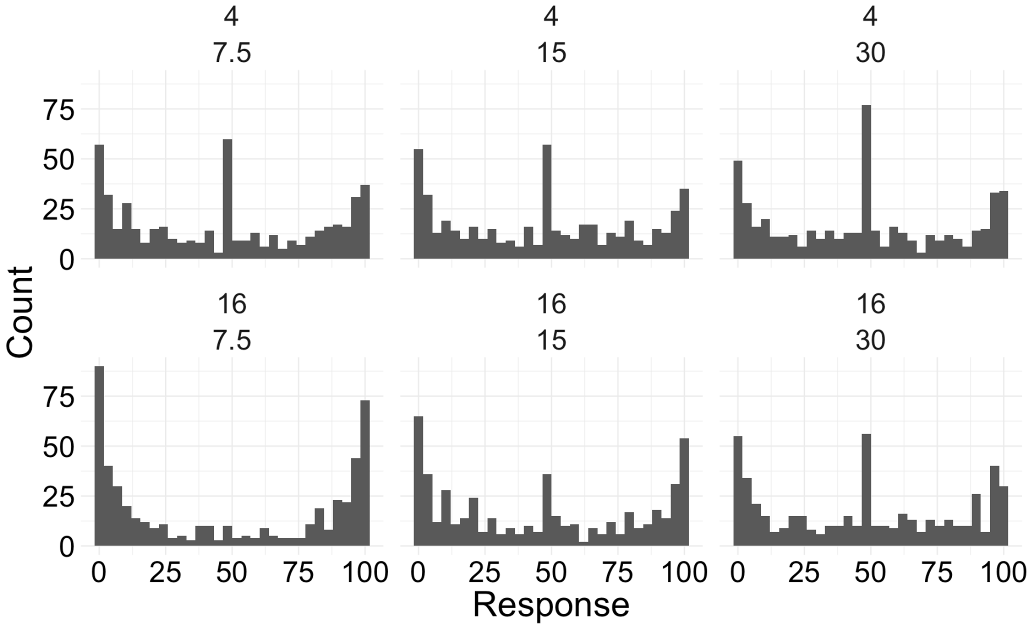


*Note*. We observed distinct peaks in confidence responses at 0 and 100, which varied as a function of sample size (4 observations in the top panel and 16 observations in the bottom panel) and standard deviation: small (left panels), medium (middle panels), and large (right panels).

**Figure S2**

Fit of Sample Mean Model to Accuracy and Confidence Data

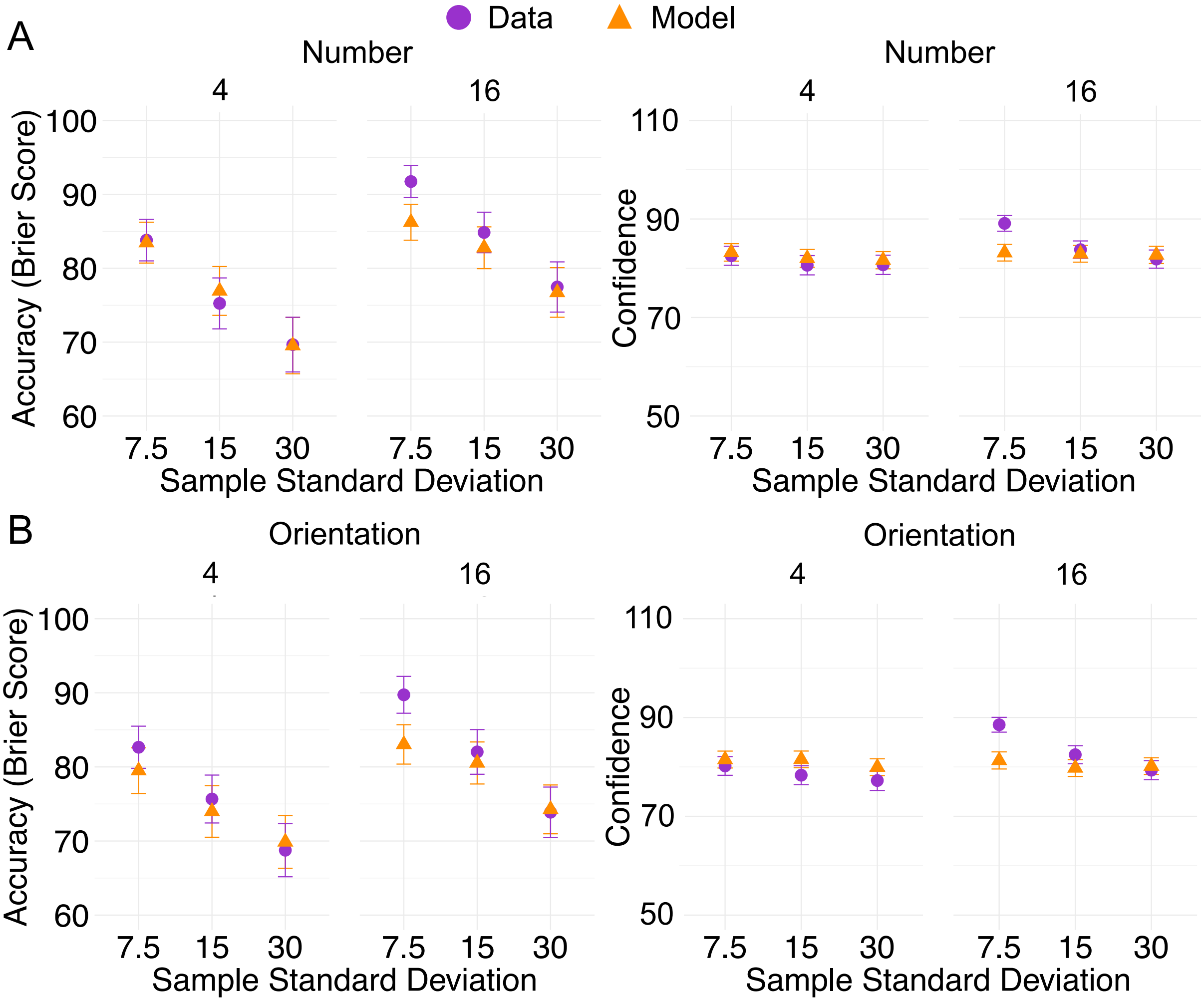


*Note*. Model predictions from the sample mean model (orange) and data (purple) for the orientation (A) and number (B) task. Error bars reflect ±1 SEM across participants within each condition. For the model, we generated a simulated dataset from the posterior predictive distribution (i.e., draws from the joint posterior over parameters and observations).

**Figure S3**

Fit of Sample Size (*n*) Model to Accuracy and Confidence Data

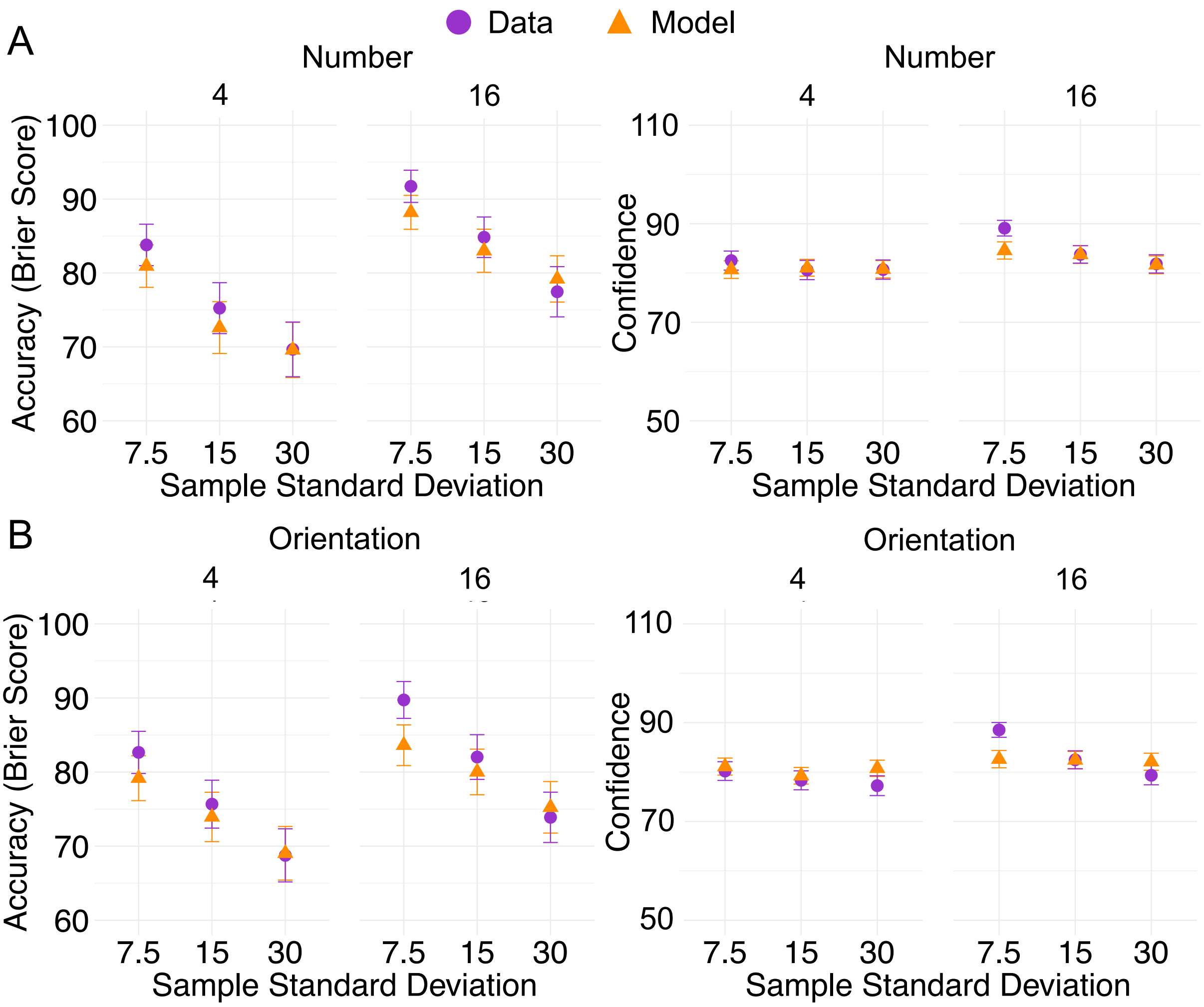


*Note*. Model predictions from the sample size model (orange) and data (purple) for the orientation (A) and number (B) task. Error bars reflect ±1 SEM across participants within each condition. For the model, we generated a simulated dataset from the posterior predictive distribution (i.e., draws from the joint posterior over parameters and observations).

**Figure S4**

Fit of Standard Deviation (*SD*) Model to Accuracy and Confidence Data

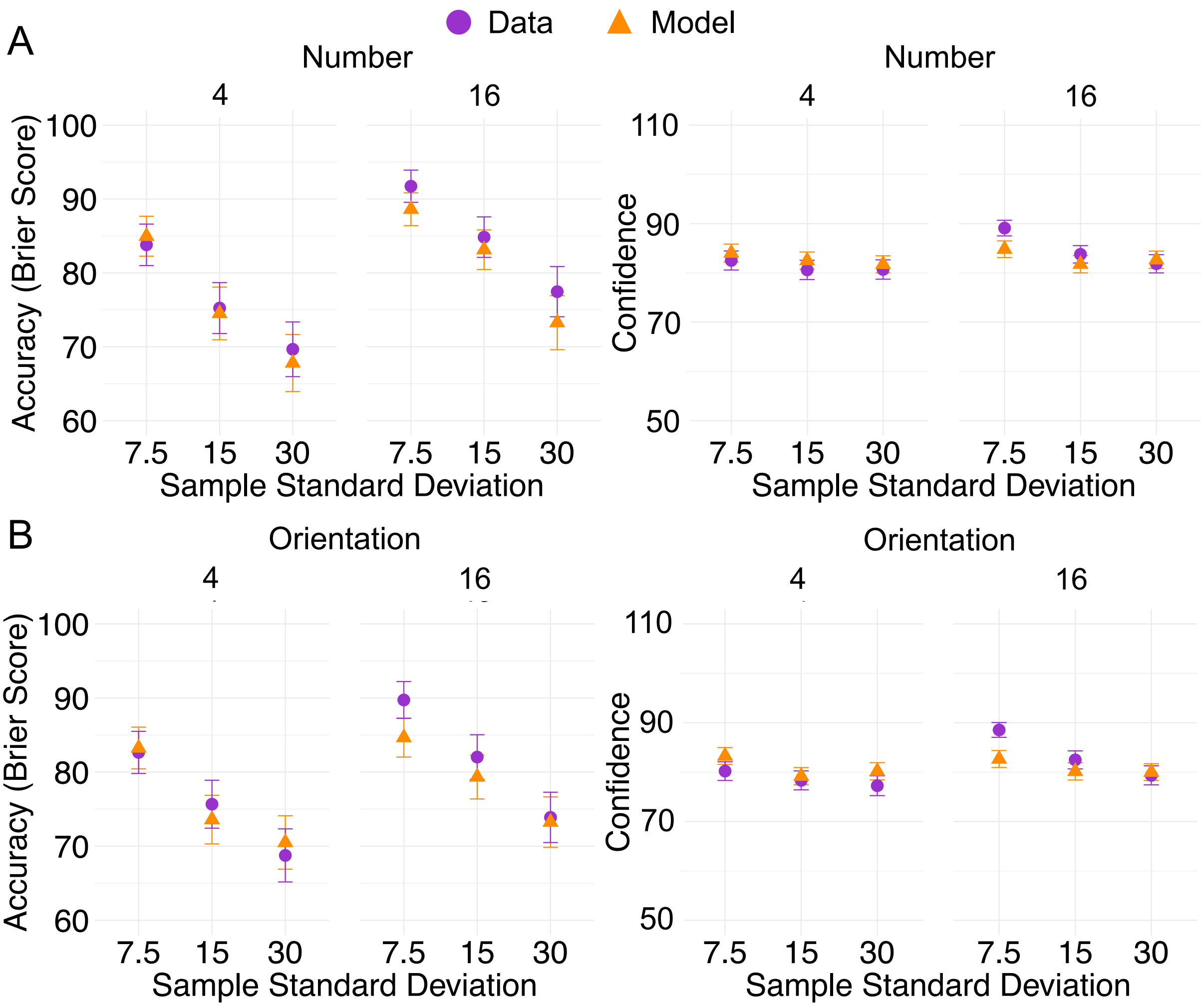


*Note*. Model predictions from the sample standard deviation model (orange) and data (purple) for the orientation (A) and number (B) task. Error bars reflect ±1 SEM across participants within each condition. For the model, we generated a simulated dataset from the posterior predictive distribution (i.e., draws from the joint posterior over parameters and observations).

**Figure S5**

Fit of Additive (SD + N) Model to Accuracy and Confidence Data

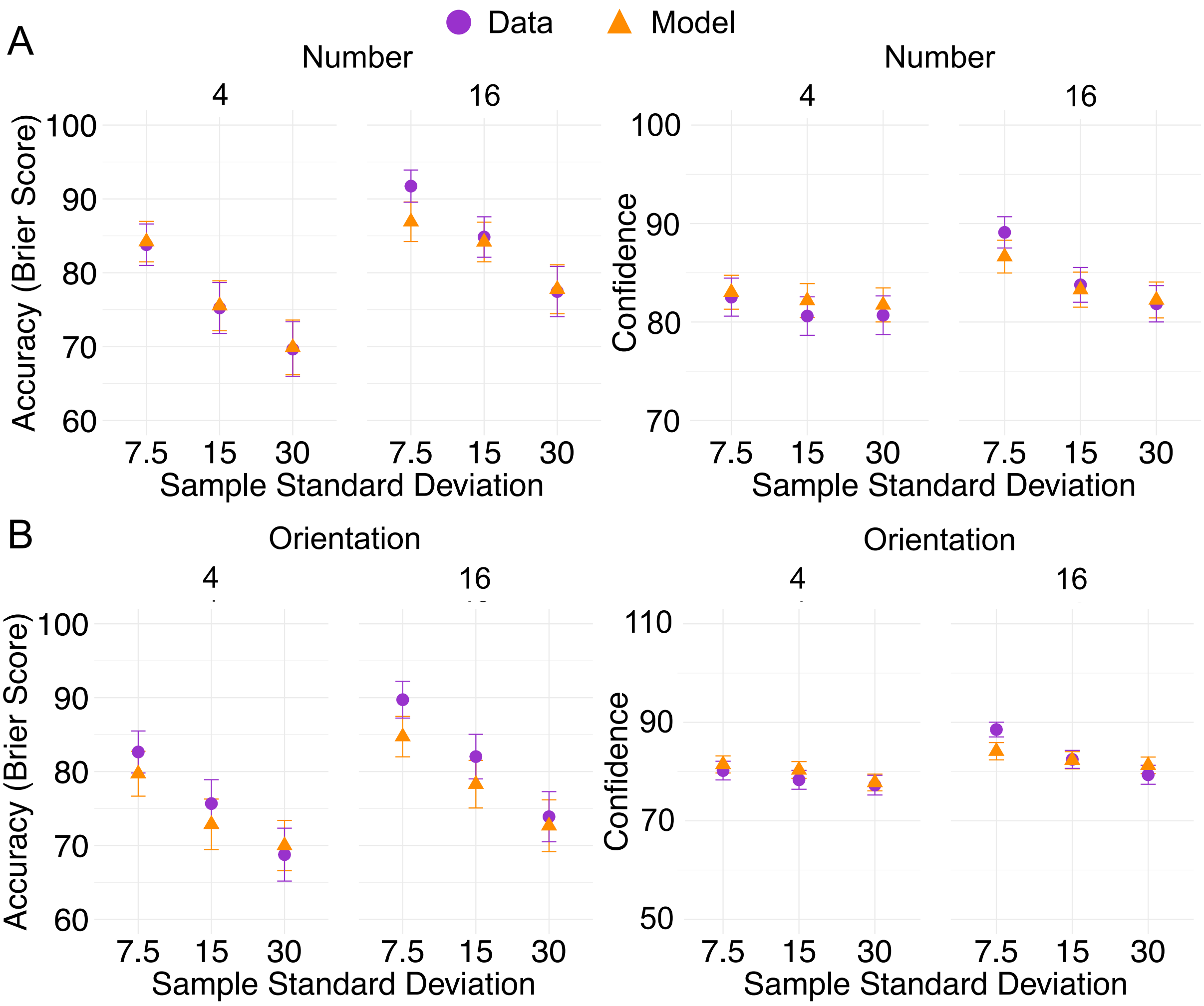


*Note*. Model predictions from the additive sample size and sample standard deviation model (orange) and data (purple) for the orientation (A) and number (B) task. Error bars reflect ±1 SEM across participants within each condition. For the model, we generated a simulated dataset from the posterior predictive distribution (i.e., draws from the joint posterior over parameters and observations).

**Figure S6**

Fit of Interactive (SD * N) Model to Accuracy and Confidence Data

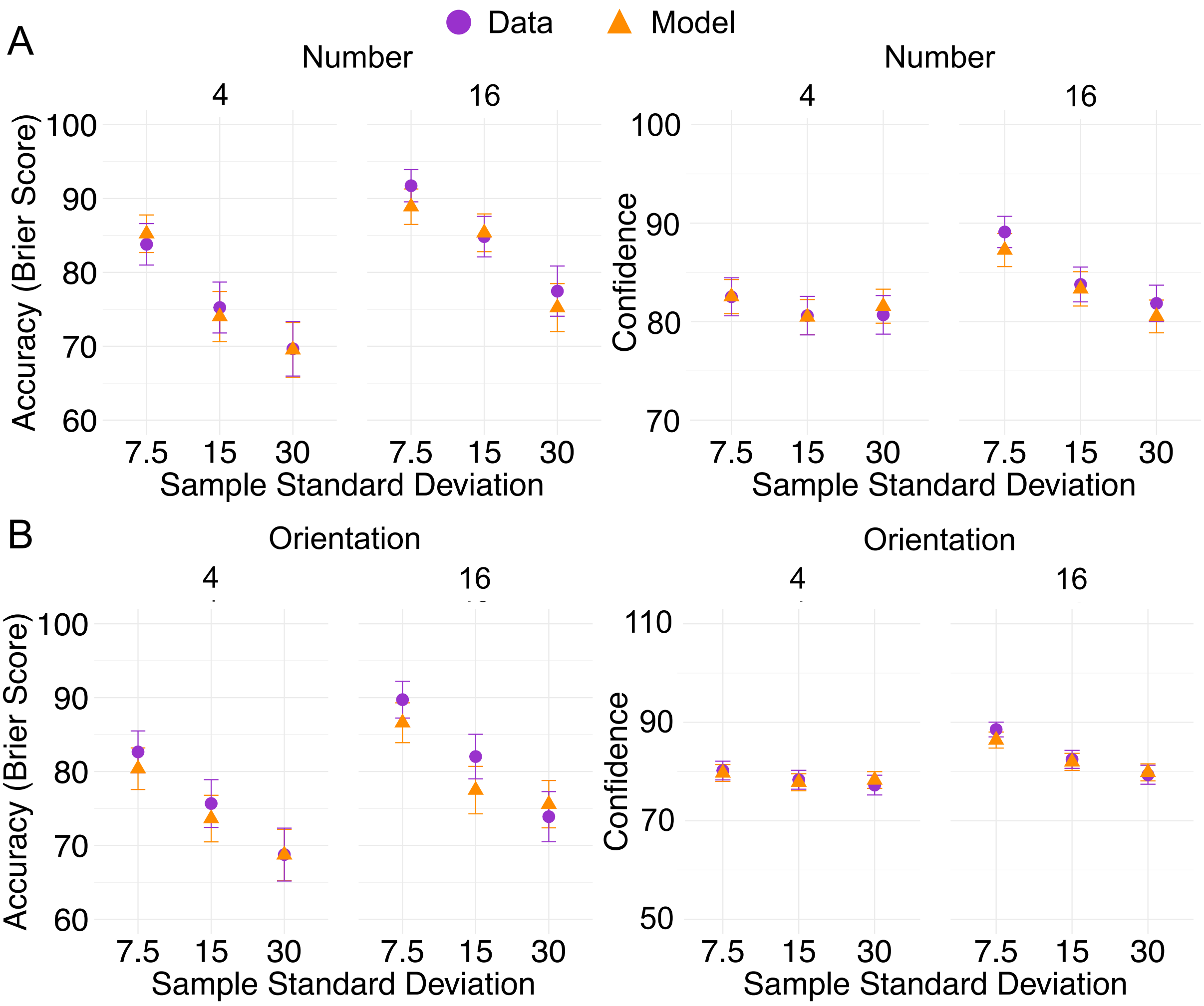


*Note*. Model predictions from the interactive sample size and sample standard deviation model (orange) and data (purple) for the orientation (A) and number (B) task. Error bars reflect ±1 SEM across participants within each condition. For the model, we generated a simulated dataset from the posterior predictive distribution (i.e., draws from the joint posterior over parameters and observations).

**Figure S7**

Fit of Bayesian Model to Accuracy and Confidence Data

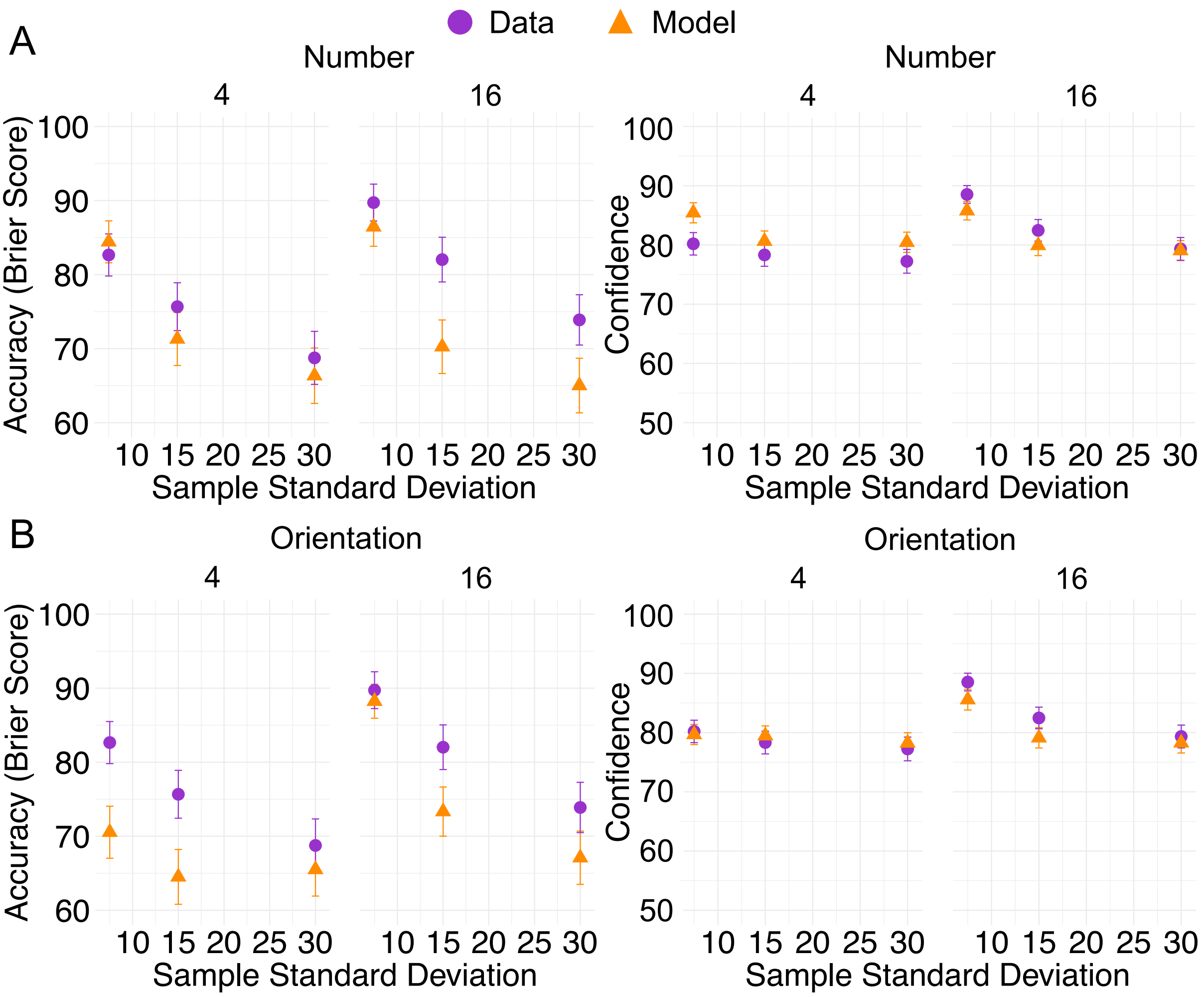


*Note*. Model predictions from the Bayesian model (orange) and data (purple) for the orientation (A) and number (B) task. Error bars reflect ±1 SEM across participants within each condition. For the model, we generated a simulated dataset from the posterior predictive distribution (i.e., draws from the joint posterior over parameters and observations).

**Figure S8**

Model and Observed Slider Responses Across Standard Error Conditions

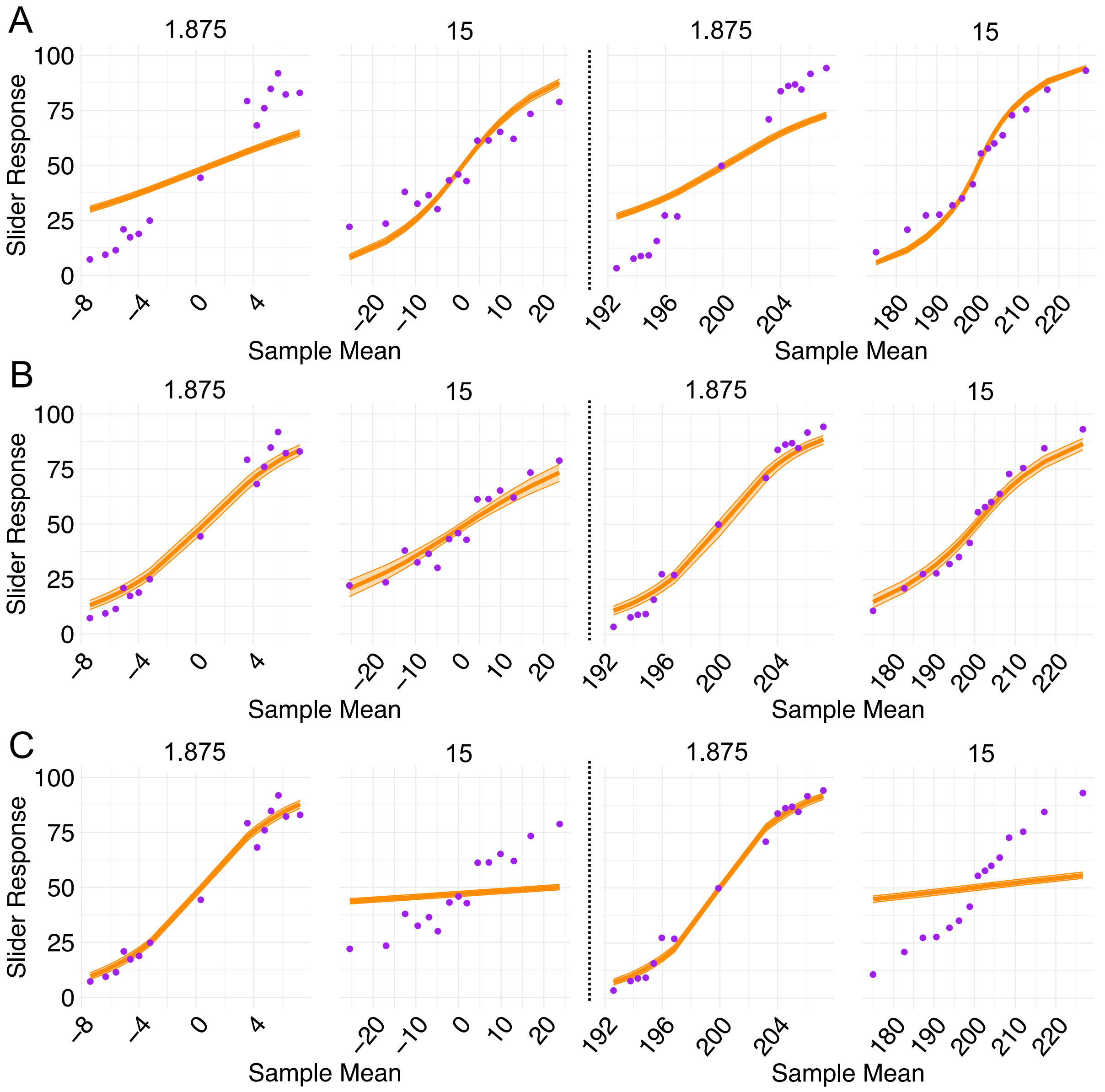


*Note*. Model predictions (orange) and empirical data (purple) for the orientation (left panels) and number (right panels) tasks. (A) The *mean-only* model, in which responses are based solely on the sample mean, fails to capture the steep increases in confidence for low–standard-error samples. (B) The *standard error* model, in which responses are based on the sample mean scaled by the standard error, successfully captures the steep confidence changes for low–standard-error samples and the shallower changes for high–standard-error samples. (C) The *Bayesian* model, in which responses are based on the posterior probability ratio, fails to capture the shallow confidence changes observed for high–standard-error samples. Error bars indicate 95% credible intervals for the model's expected values, computed from posterior samples.

**Figure S9**

*Comparing Domain-Specific vs. Domain-General Parameter Settings*

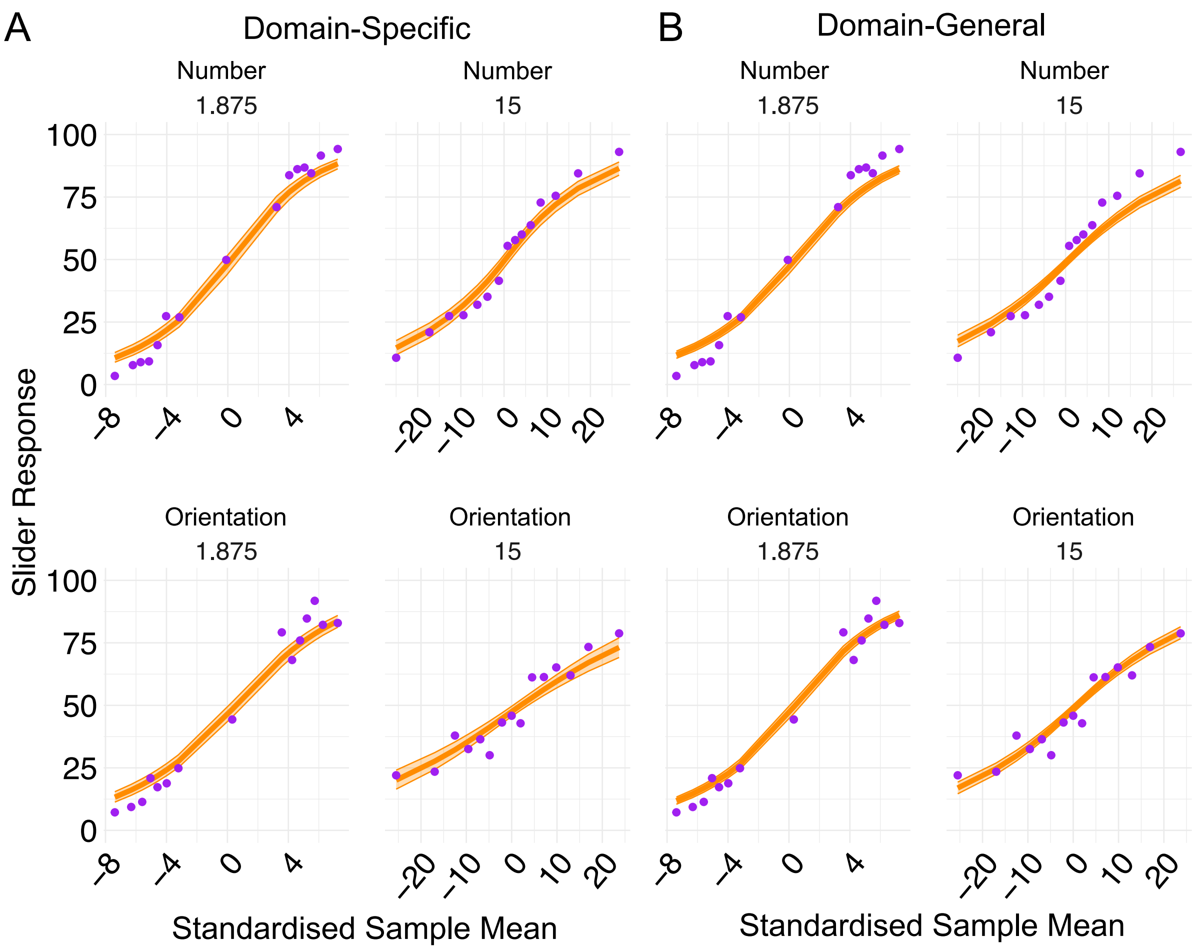


*Note.* Model predictions (orange) and empirical data (purple) for the number (top panels) and orientation (bottom panels) tasks. (A) The *domain-specific* model used the standard error framework with separate parameter estimates for each task. (B) The *domain-general* model used the same framework with shared parameter estimates across tasks.
